# Bridging Viruses and Prokaryotic Host through Miniature Inverted-repeat Transposable Elements (MITEs)

**DOI:** 10.1101/2024.01.18.576219

**Authors:** Francisco Nadal-Molero, Riccardo Roselli, Silvia Garcia-Juan, Alicia Campos-Lopez, Ana-Belen Martin-Cuadrado

## Abstract

Transposable elements (TEs) have a pivotal role in the evolution of genomes across all life domains. “Miniature Inverted-repeat Transposable-Elements” (MITEs) are non-autonomous TEs mainly located in intergenic regions, relying on external transposases for mobilization. The boundaries of MITEs’ mobilome were explored across nearly 1700 prokaryotic genera, 183232 genomes, revealing a widespread distribution. MITEs were identified in 56.5% of genomes, totaling over 1.4 million cMITEs (cellular). Cluster analysis revealed that a significant 97.4% of cMITEs were conserved within genera boundaries, with up to 23% being species-specific. Subsequently, this genus-specificity was evaluated as a tool to link microbial host to their viruses. A total of 51655 cMITEs had counterparts in viral sequences, termed vMITE (viral), resulting in the identification of 2798 viral sequences with vMITEs. Among these, 1501 sequences were positively assigned to a previously known host (41.8% were isolated virus, and 12.3% were assigned through CRISPR data), while 379 new host-virus associations were predicted. Deeper analysis in Neisseria and Bacteroidetes groups allowed the association of 242 and 530 new additional viral sequences, respectively. Given the abundance of non-culturable virus sequences accumulated in databases lacking affiliations with their microbial targets, MITEs are proposed as a novel approach to establishing valid virus-host relationships.

**GRAPHICAL ABSTRACT:** 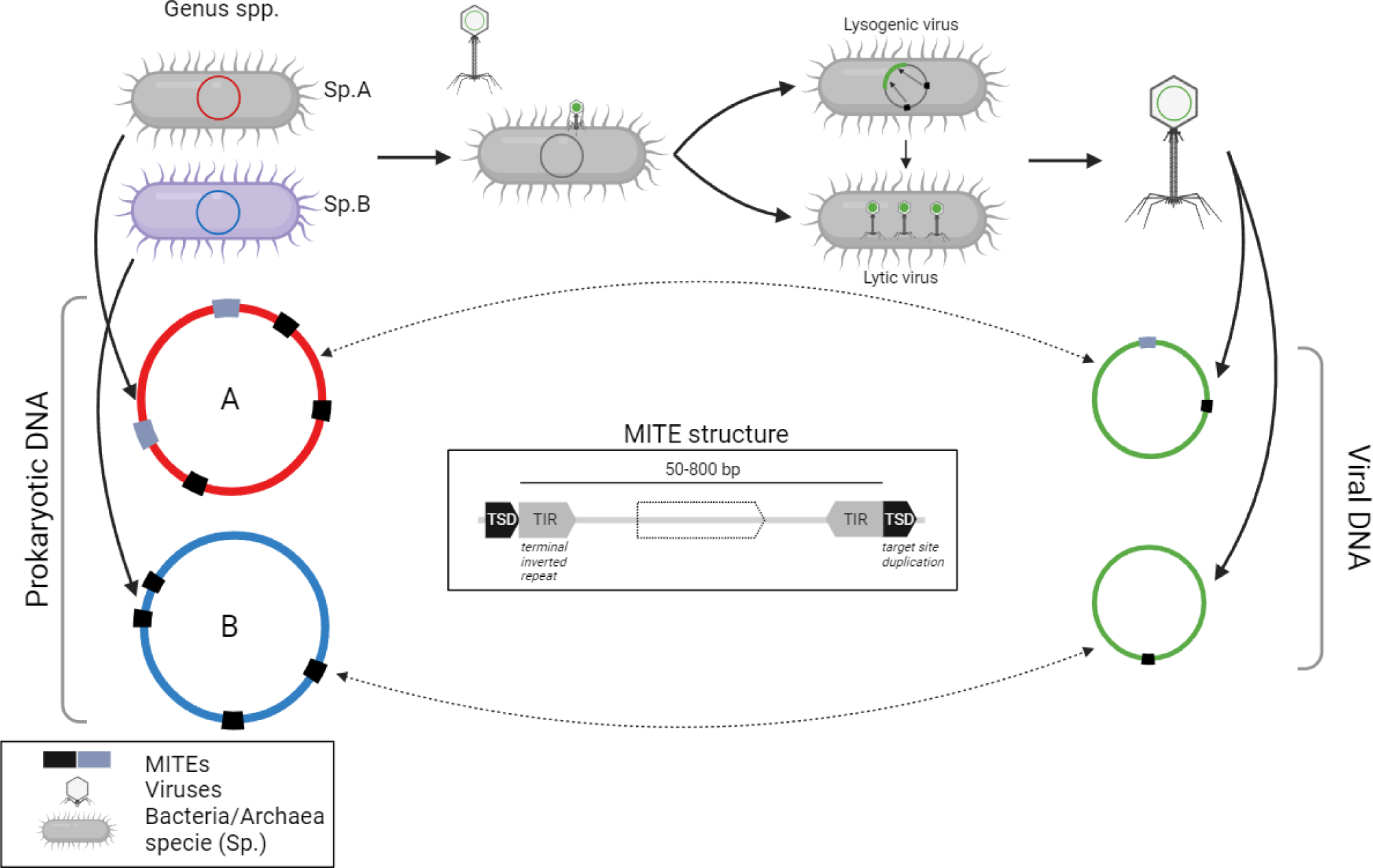

## INTRODUCTION

Prokaryotic cells and their viral predators have been evolving together for billions of years (1,2). The identification of the microbial host of many environmental viruses, mainly known by their assembled sequence, is one of the Microbiology Gordian knots. Nowadays, further than the virus isolation from a culturable host, the most reliable tool for identifying its potential host involves the presence of viral proto-spacers in the CRISPR-Cas arrays (3,4), a system found in 50% of the bacterial genomes and nearly 90% of the archaeal ones (5). Other approximations rely in genomic features such as *k-mers* (used in the bioinformatics programs WIsH (6), VHM-net (7) and PHP (8)), different oligonucleotide frequencies (9–12), or similar codon use (Labonté et al., 2015). Also, the identification of exogenous DNA within a genome as an interchanged DNA fragment swapped at the infection stage, could be also very useful for their association; e.g., tRNAs (13), bacterial genes within the virus genome (14) or similar protein content (15,16).

Along with the interchanged genomic material among viruses and hosts, there is also the possibility to shift self-transmissible mobile genetic elements such as insertion sequences (IS), transposons or integrons (17). These MGEs play a pivotal role as recombination hot-spots that promote horizontal gene transfer (HGT) events (18) and, in fact, the genomic mosaicism often observed in bacteriophage genomes has been suggested to be the result of these rearrangements (19,20). However, despite the multiple HGT events described between viruses and their hosts, the numbers of transposable elements (TEs) found in viral genomes are low (21), and for example, IS elements, which are the most frequent class of prokaryotic TEs, have been found frequently in plasmids but rarely in phages, reflecting a very strong and efficient post-insertional purifying selection. Meanwhile prokaryotes constantly exchange genetic material trespassing inter-species barriers that impact their genome evolution, TEs contribution to viral genome evolution remains largely unexplored. Previously, Zhang et al (2018), detected the presence of specific TEs called ‘Miniature Inverted-repeat Transposable Elements’ (MITEs) in several eukaryotic viral genomes from Ascaoviriae, Polydnaviridae and Pandoraviridae families (23,24), and although no equal MITEs were found in their hosts, similar sequences were found and a link among viruses and hosts might be stablished based on their similarity (64-97%).

MITEs are considered non-coding sequences of small size (50-800 bp) (Figure 1) that include an internal DNA sequence flanked by terminal inverted repeats (TIR) of at least 10-15 bp (25). Target Duplication Sites (TSD) of 2-10 bp flank the MITE sequence as a result of the MITE insertion into the DNA target site; however, these TSD not always are found in prokaryotes MITEs (23). Although not totally clear, MITEs are thought to be derived from internal deletions occurred at ISs, leaving the terminal repetitions as traces of the ancestral transposition events (26,27). Importantly, MITEs are classified as a class II non-autonomous transposable elements due to the lack of transposase and the need to use other MGEś transposase to promote their transposition (28). Then, given the complete dependency from external enzymes to propagate themselves across genomes, MITEs could be considered *de facto* parasites of other MGEs; furthermore, “double parasites” in the case of viral MITEs.

**Figure 1.**
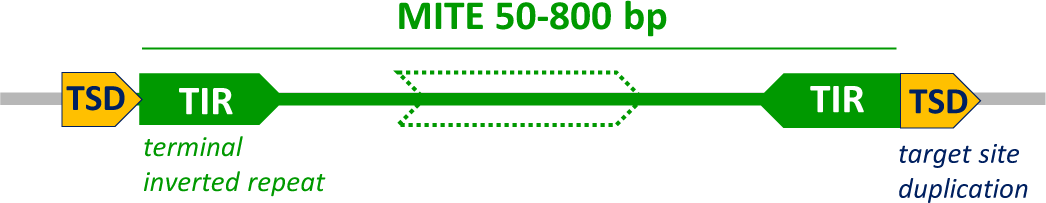
Structure of a MITE element, which comprises two inverted repeat sequences (TIR in green) at both ends (50-800 pb) limited by flanking regions of equal target sites (TSD in yellow).

MITEs number varies among dozens in specific eukaryotic virus (22), hundreds in microbial genomes (30), or up to thousands copies in eukaryotic cells (29). They have been described to play a significant role in the evolution of several eukaryotes through the increase of genome size, formation of new genes and regulation of gene expression (23,31–35). However, their roles are much less known in prokaryotes. The first MITEs described in Bacteria were the so called ‘Correia elements’ of *N. gonorrhoeae* and *N. meningitidis* (36,37), and their destabilizing role of the genome was demonstrated after the analysis of the transcriptional regulation of several *Neisseria* genes (38). Negative effects related to transposition events have been described so far, such as gene inactivation, alteration of gene function, genomic rearrangements, modification of the genome structure (formation of secondary structures), or genomic instability leading to accumulation of mutations (30,39–48)). Importantly, MITEs have been also related with the genes dispersion of antimicrobial resistance in *Acinetobacter* and *Enterobacter cloacae* (49–52). Despite their presence in multiple prokaryote genomes (30), little investigations have been performed so far further than in some *Neisseria* (53,54) and *Sulfolobus* strains (45,55).

In this work, it is shown that a surprisingly vast and previously untapped diversity of MITEs are present in bacterial and archaeal genomes. Also, but in less degree, in viral sequences. The taxon-specificity observed among prokaryotic MITEs suggests that the presence of the same MITE sequence in both viral and prokaryotic genome could serve as a tool to associate a virus with its putative hosts. It is also discussed the dynamics of MITEs across and within species in the context of their taxonomic specificity.

## MATERIALS AND METHODS

### Prokaryotic and viral Sequences

A total of 8220 archaeal genomes were downloaded from the Assembly database at NCBI from its GenBank collection. For Bacteria, the RefSeq collection was downloaded from Assembly database in NCBI, n=175 012 genomes (available in August 2022). A total of 183 232 prokaryotic genomes, were used in this study. Viral sequences were obtained from the GenBank database of the NCBI (ftp.ncbi.nlm.nih.gov/genomes/Viruses/AllNucleotide/AllNucleotide.fa, n=10453098, cointaining vrial genomes and gene-amplicons, December 23, 2021) and from the IMG/VR v.4.1 database (n=5621398 sequences, (56)). Human feces metaviromes from healthy adults samples used in Supplementary Table S10 were obtained from the following projects; PRJNA491626 (SRR7892428 and SRR7892426) (57); PRJEB9524 (ERR906942 and ERR906934) (58) and PRJNA318788 (SRR3403840) (59). Crassphages already published were also used as references in the analysis.

### MITE Detection

The program used for the detection of MITEs was MITE Tracker (60) with default search parameters. To ensure that MITEs detected were *bonafide* non-autonomous elements, those sequences containing complete or partial transposases were removed. For this purpose, 815 857 transposases were downloaded from Uniprot and *nr* database from NCBI and were used in a BlastX comparison (*cut-off:* 30% of amino acid similarity). A total of 935 338 sequences were excluded. The final collection consisted in 1 406 057 MITEs of Bacteria, 18 091 of Archaea and 1 726 from virus. Cellular MITEs were named cMITEs and, viral ones, vMITEs. Similar MITEs detected in viral sequences, but containing miss-matches *versus* de cMITEs, the so called *pseudo*-vMITEs (*ps*-vMITEs), were detected by BlastN comparisons using cMITEs as a query *versus* the viral databases (*cut-off*: 95% id in 100% of their length). A total of 5235 *ps*-vMITEs were identified.

### Clustering

Cd-hit-*est* program was used to cluster the MITEs using a *cut-off* of 95% id in 100% of the sequence length (-aS 1) (61). Cluster representatives were considered as unique MITEs. In order to minimize the number of cMITEs to represent, bacterial cMITEs were clustered at 100% id in their complete length and representative sequences were used to reconstruct the net.

### Bioinformatic analysis of MITEs

GC-content from the bacterial and archaeal genomes was calculated using the SeqTk v.1.3-r106 package (https://github.com/lh3/seqtk.git). Prodigal.v2.6.3 (62) was used in order to define the ORFs of the IMG/VR v.4.1 viral sequences. The coding density of the genomes was calculated dividing the length of the ORFs by the length of the complete genome. To assess their intra-gen or inter-gen positions, the ‘start’ and ‘end’ MITE coordinates given by MITE Tracker were used to evaluate their locations across the genomes. The taxonomy associated to each MITE was that of the associated genome accordingly the NCBI taxonomy classification. Putative MAGs contamination was calculated with CheckM v1.1.3 (63). ISFinder database was used to compare MITE sequences extracted with previous IS elements already described (64). Prophages prediction was done with CheckV (65)). Accession numbers of sequences of the inlet of Figure 6 and the Supplementary Figure S7 are: 1-BK042455, 2-BK045475, 3-UViG-2541047207, 4-UViG-2541047202, 5-UViG-2541047209, 6-UViG-2547132244, 7-UViG-2744054866, 8-UViG-2541047237, 9-UViG-644736394, 10-UViG-3300008695-1, 11-BK043181, 12-BK056678, 13-UViG-3300008695-2, 14-BK016719, 15-UViG-3300008746, 16-UViG-3300007335, 17-UViG-3300008140, 18-UViG-3300007294, 19-BK058633, 20-UViG-3300008695-3, 21-UViG-7000000395, 22-UViG-2639762804, 23-UViG-2541047239, 24-UViG-2541047182, 25-UViG-2541047193, 26-BK041853.

### Statistical analysis

Statistical analyses were conducted in R version 4.2.1 (Team RC. R: A language and environment for statistical computing. Vienna: R Foundation for Statistical Computing; 2019) and RStudio version 2022.07.0 (Team R. RStudio: integrated development environment for R. Boston: RStudio, PBC; 2022.). Pearson correlations were calculated using *cor.test* function of the stats (v. 3.6.2) R package for numeric variables of paired samples at 0.95 of confidence level.

### Viral Nets

Bacterial and archaeal cMITEs were calculated with SSNetwork (66)) at 95% id in 99% of the sequence length. vConTACT2 0.9.09 was used to classify the viral contigs containing MITEs using the Prokaryotic Viral RefSeq (version 221 with ICTV and NCBI taxonomies). Only those results with a score>5 were considered. The nets were visualized with Cytoscape 3.10.1 (67).

## RESULTS

### Census of cMITEs in Prokaryotic RefSeq genomes

The microbial cellular MITEs (cMITEs) were identified in 183 232 Ref-Seq microbial genomes (Bacteria and Archaea) using the program MITE-Tracker (60). This analysis, which comprises over 400 families and nearly 1700 prokaryotic genus, revealed more than one million of cMITEs sequences unequally distributed among bacterial and archaeal (Table 1), being present in 58.5% of the bacterial genomes (1 406 057 cMITES (111 244 unique) in 102 498 genomes, Supplementary File S1) and in 20% of the archaeal ones (18091 cMITEs (7114 unique) in 1015 genomes, Supplementary File S2). BlastN comparisons against the ISFinder database showed 1.6% of the cMITEs to be similar to MITEs previously described, mainly with MITEEc1_IS630 (49.31%), IS1106_IS5 (24.5%) and ISNme5_IS110 (21.7%), present in *N. meningitidis* and *Escherichia coli*. Interestingly, among the genomes hosting cMITEs, only 2.5% had one single cMITE, while 91.7% harbored an average of 11 different cMITEs.

**Table 1.** Main genomic features of the MITEs found in Bacteria, Archaea and Viral sequences.

|  |  | Archaea | Bacteria | Virus GB | Virus JIMG/VR |
| --- | --- | --- | --- | --- | --- |
|  | # genomes/sequences analyzed | 8220 | 175012 | 10468678 | 2377994 |
|  | # genomes/sequences with MITEs | 1015 | 102498 | 361 | 310 |
| MITEs features | # MITEs | 18091 | 1406057 | 610 | 1116 |
|  | # unique MITEs | 7114 | 111244 | 230 | 495 |
|  | minimum size (bp) | 41 | 49 | 53 | 36 |
|  | Median size (bp) | 104 | 104 | 72 | 82 |
|  | maximum size (bp) | 797 | 800 | 775 | 789 |
|  | # clusters with > 1 MITE | 7114 | 111244 | 230 | 495 |
|  | # clusters with > 2 MITEs | 2623 | 50482 | 136 | 160 |
|  | # clusters with > 5 MITEs | 949 | 23481 | 33 | 54 |
|  | # clusters with > 10 MITEs | 291 | 10288 | 11 | 14 |
|  | # clusters with > 50 MITEs | 20 | 2094 | 0 | 1 |

cMITEs were found to be independent of several genomic traits. The higher number of cMITEs or diversity (unique cMITEs) was not found in the largest genomes, and instead, two peaks were observed for 2.5 Mb and 5 Mb genomes (Supplementary Figure S1A). Also, it was investigated whether smaller cMITEs might propagate more easily or persist for longer periods in the genomes, but results showed that cMITE size and abundance were unrelated (Supplementary Figure S1B). As cMITEs are found in AT-enriched genome regions (68), it was calculated their frequency regarding the genome GC-content where they were found (Supplementary Figure S2A). The global analysis showed that low-GC genomes (30-39%) exhibited higher number of cMITEs (despite the bias in the GC-content range of the number of genomes analyzed). Importantly, a positive correlation was observed between the GC-content of the cMITE and the GC-content of the genome containing it (Supplementary Figure S2C), suggesting that mobility of cMITEs is constrained within certain genomic boundaries and might be conditioned to similar GC-content genomes. Alternatively, it could also be interpreted as a rapid adaptation by the cMITE sequence to its hosts genome. Regarding the genome coding density, most of the genomes enriched in cMITEs (n>100) exhibited coding densities between 80-85% in bacteria, and slightly higher in archaea, reallocating and maintaining a higher number of MITEs insertions (Supplementary Figure S3). Of particular interest was the cluster formed by genomes of the Spirochaetes Phylum (mainly *Leptospira*, *Sediminispirochaeta* and *Treponema*), with coding densities lower than 80% and an average of 145.4 cMITES per genome. As previously observed, majority of cMITEs were located in intergenic positions, accounting for 90.4% and 88.9% in Bacteria and Archaea, respectively (Supplementary Figure S4).

### Distribution of cMITEs across Taxonomic Ranks

Among the different species, the best represented microbes containing cMITEs were *Vibrio cholera* cMITEs, *Neisseria meningitidis* and *Streptococcus pneumoniae* (Figure 2), mainly due to the bias in the RefSeq database, with some clinical isolates in high abundance and an uneven sampling of the tree of life. Results showed that, for 28.7% of the microbial species, 100% of their strains hosted cMITEs in their genomes. Conversely, also for a similar number of species, 30.5%, only 10% of the strains (or fewer) had cMITEs in their genomes (note the “U-shaped” graph in Supplementary Figure S5). These findings, together those state ahead in this section, may indicate that once a cMITE is settled in a genome and in the absence of systems that control their proliferation, the possibility of spreading within its host genome and also migrate to the genome of a related species genome, it is quite high. This trend, microbial species having cMITEs in most the genomes sequenced or none at all (and little falling in between) was observed when analyzed genera with more than 50 genomes available (Figure 3). The presence of cMITEs within a genus with more than 1000 genomes available was pretty remarkable and, for example, 99.8% of the genomes of *Shigella* contained a cMITE, 99.6% in the case of *Legionella*, 99.4% of *Neisseria*, 96.4% of *Salmonella* or 88.8% in the case of *Escherichia* genomes (Supplementary Table S1). Meanwhile, a strong resilience to acquire these MGEs was observed for *Mycobacteroides* genomes (0.2%), *Staphylococcus* (0.4%), *Helicobacter* (1.1%), *Klebsiella* (3.8%), *Acinetobacter* (7.3%) or *Enterococcus* (8%). When this analysis was repeated but using all the available genomes for some clinically important genera (nor just the RefSeq), such as *Neisseria* or *Staphylococcus*, a similar a trend was obtained. Specifically, over 94% of the 5837 Neisseriales genomes contained cMITEs, while persistently, they were found in only 0.01% of the 90280 *Staphylococcus* genomes analyzed (Supplementary Table S2). Complete screenings using other ecological relevant groups, such as 1064 genomes of *Pelagibacter* or the 1184 of *Prochlorococcus,* reverted zero cMITEs among their genomes.

**Figure 2.**
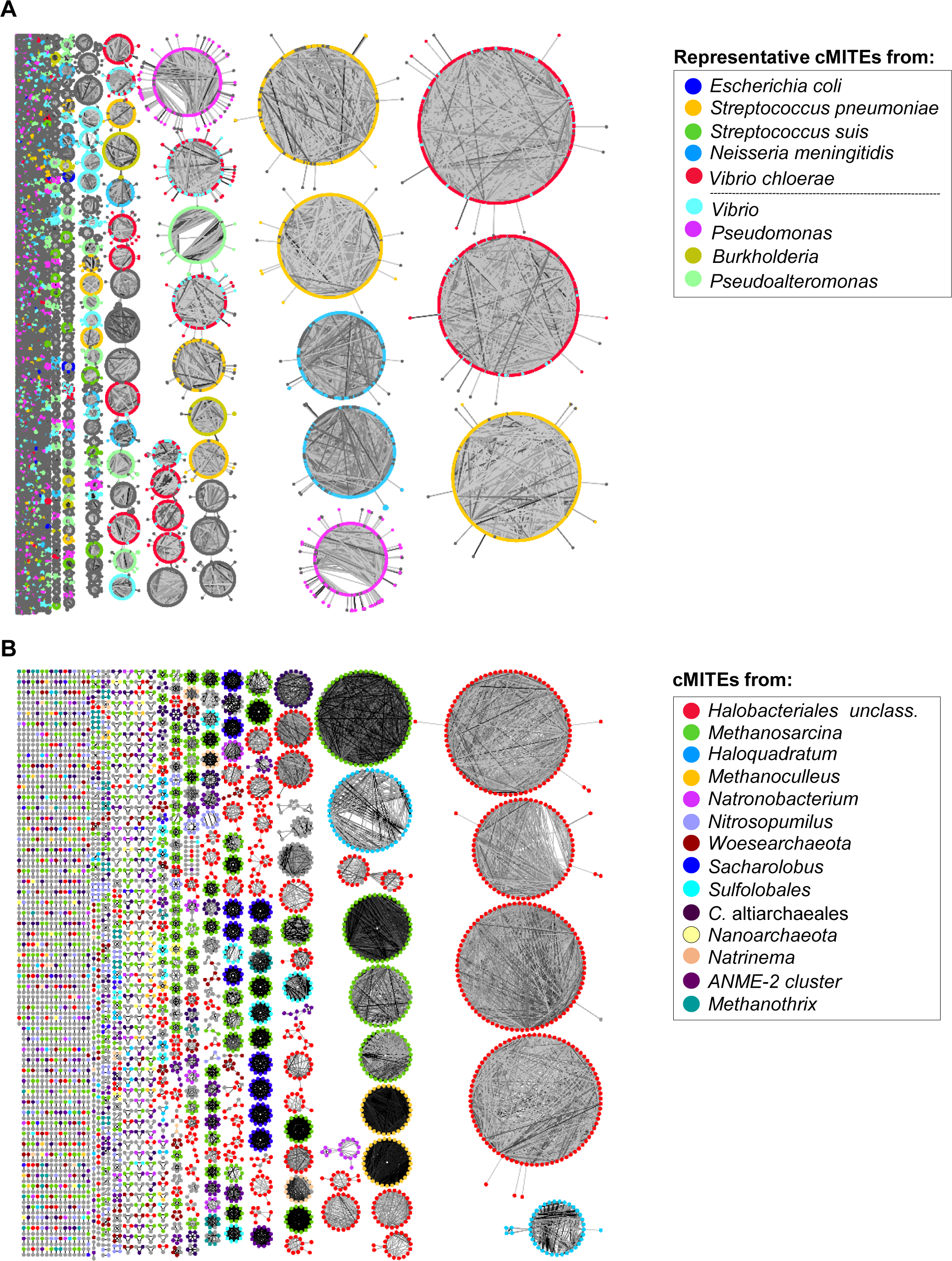
**(A)** Relations among the representatives of Bacterial cMITEs (*cuf off*: 100% id). **(B)** Relations among all the archaeal cMITEs. Both nets were constructed using an identity of 95% in 99% on the length of the cMITE sequence using SSNetwork and represented with Cytoscape (v. 3.10.1). (Only major groups were colored).

**Figure 3.**
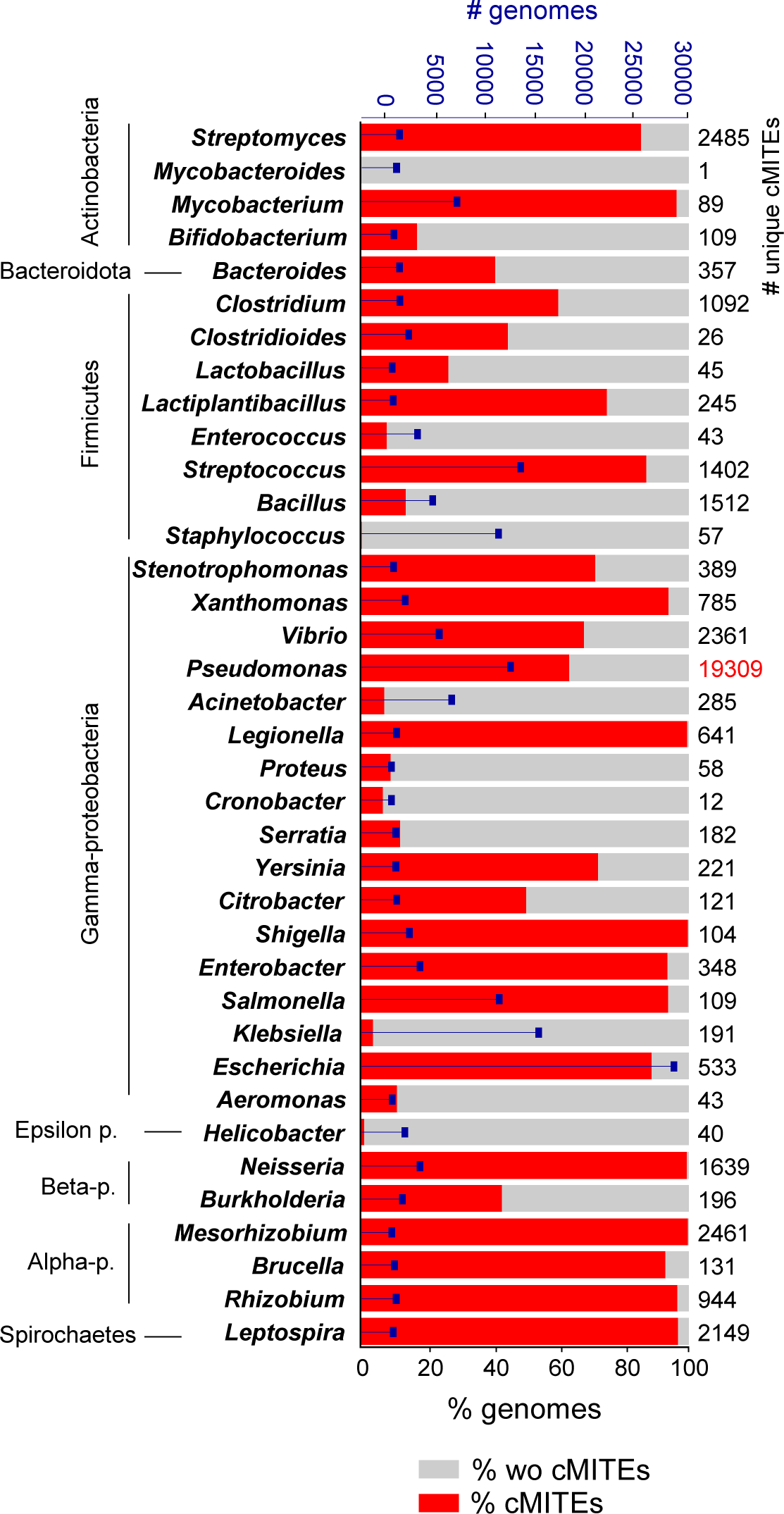
Bars represent the percentage of genomes within a genus which have cMITEs in their sequences (only those genera with >50 sequenced genomes were considered). Blue squares indicate the number of total genomes sequenced for that particular genus (upper *x axe*). Numbers of unique cMITEs is indicates on the right.

At the order level, Leptospirales, Cyanobacteria, Neisseriales and Alteromonadales genomes showed to be particularly enriched (media of 153.66, 55.31, 46.58 and 27 MITEs/genome, respectively) (Figure 4A, Supplementary Figure S6A and Supplementary Table S3). The highest number of cMITEs was found in the filamentous Cyanobacteria *Prochlorothrix hollandica* PCC9006, with 468 MITES (161 unique), followed by *Aphanizomenon flos*-*aquae* 2012/KM1/D3 and two *Parashewanella spongiae*. Within the Archaea, the *Halococcaceae* family presented the higher number of cMITEs per genome, but it was an unclassified *Haloarculaceae* (GCA_003021305.1) the one with more cMITEs, 251. This was followed by *Methanosarcina spelaei* and *Halovenus* sp003021015 (Figure 4B, Supplementary Figure S6B and Supplementary Table S3). Considering the cMITEs density (Supplementary Table S4), genomes of endosymbionts *Wolbachia pipientis* (123.4 cMITEs/Mb) and two Rickettsias had the highest values and up to 1.23% of their genomes would be constituted by cMITEs. Among Archaea, the DPANN genomes of Candidatus *Altiarchaeum hamiconexum*, a *Woesearchaea* and a strain of *Halobacteriales*, presented the highest values, 145.5, 105.1 and 99.7 MITEs/Mb, respectively. Regarding the diversity of cMITEs among taxa, it exhibited a considerable variability (Supplementary Table S4) and, for instance, *Pseudomonas* genera contained more than 19000 different cMITEs, while others as *Citrobacter* contained one single cMITE. The highest diversity was found also in two Cyanobacteria, *Okeania* sp. KiyG1 and *Okeania hirsute,* with 225 and 221 different cMITEs from a total of 278 and of 320, respectively (followed by *Companilactobacillus heilongjiangensis* (211/241) and *Thiocapsa rosea* (202/ 294)).

**Figure 4.**
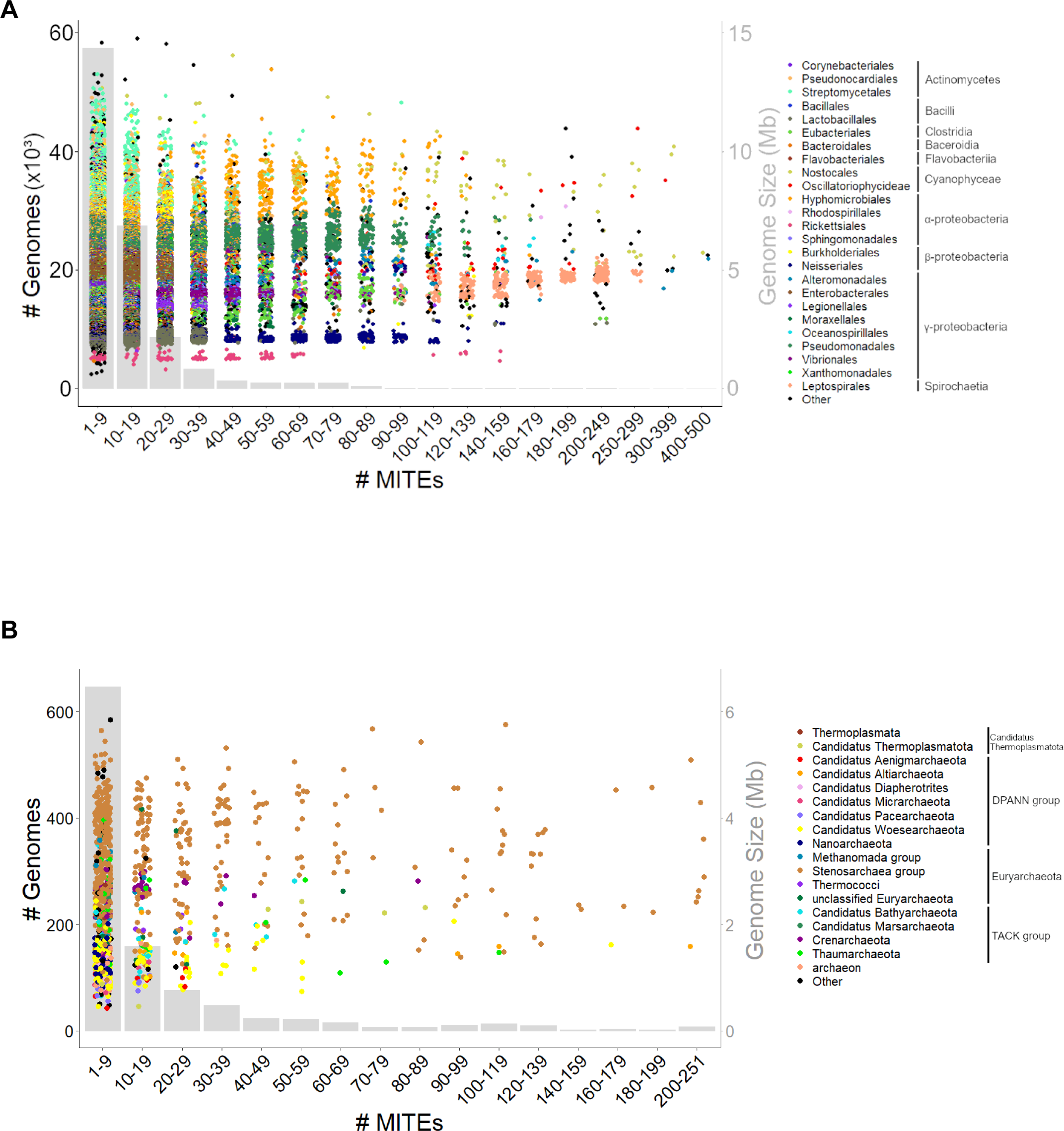
Distribution of MITEs sequences among bacterial (A) and archaeal (B) genomes accordingly with their taxonomy and number of cMITEs. Grey bars indicate the number of genomes classified accordingly with the cMITEs number. Each dot represents a genome colored accordingly its taxon (order). Right-*y* axe indicates its genome size.

### Specificity of cMITEs

Mobility boundaries of cMITEs were inspected to know whether hosting a specific cMITE sequence was a common trend maintained by phylogenetically related members, or also shared across distant taxonomic ranks. Clustering analysis led to a total of 118359 clusters of cMITEs (*cut-off*: 95% id, 100% length coverage) with a striking 98.2% of these clusters constituted by cMITEs that belonged to genomes from the same genus, a high specificity for a mobile element (Table 2). Near 23% of the cMITEs resulted to be species-specific, and independently of the number of repetitions, it could be only found in one single prokaryotic species. To visualize the specificity, a net was constructed with de-replicated cMITEs representatives of archaea and bacteria (Figure 3), and regardless the variability of cMITEs intra-genus, each circle was formed mainly by cMITEs shared among the genomes of one single specie or genera and not beyond. The average of genomes sharing the same cMITE was 8.9, but this number may increase up to 17341 such as the case of *Escherichia* or *Shigella* genomes.

**Table 2.** Specificity of cMITEs by taxon rank. Distribution of bacterial and archaeal cMITEs in clusters (*cut-off*: 95% identity in the complete length). According to the taxon of each cMITE, clusters were classified as mixed if cMITEs from different genera, family, order, class of phylum were found in the same cluster.

|  | **Archaea** | **Bacteria** |
| --- | --- | --- |
| **# cMITes** | 18091 | 1406057 |
| # cMITes species-specific | 14592 | 322157 |
| # cMITes genus-specific | 16189 | 1369639 |
| **# cMITes of mixed clusters** | 926 | 67975 |
| # cMITes of mixed-genus clusters | 412 | 29013 |
| # cMITes of mixed-family clusters | 235 | 22865 |
| # cMITes of mixed-order clusters | 9 | 484 |
| # cMITes of mixed-class clusters | 10 | 6613 |
| # cMITes of mixed-phylum clusters | 260 | 0 |
| **# clusters** | 7114 | 111245 |
| # clusters species-specific | 6665 | 90492 |
| # clusters genus-specifics | 6937 | 109343 |
| # clusters of mixed | 70 | 1814 |
| # clusters of mixed-genus | 44 | 1085 |
| # clusters of mixed-family | 7 | 693 |
| # clusters of mixed-order | 2 | 27 |
| # clusters of mixed-class | 2 | 9 |
| # clusters of mixed-phylum | 15 | 0 |

However, exceptions of the specificity were found in 2192 mix-clusters, counting 436464 cMITEs (see, for example, the case of *Bacteroides* and *Phocaecicola* in Figure 5B). A closer inspection of these clusters could reduce this number to 1884 clusters, a 4.8% of the total number (68 900 cMITEs). Two big mixed-clusters contained one single cMITE sequence that was differently taxa-associated from a pool over 10000 sequences (these implied 75634 cMITEs, a 5.3%). Also importantly, 162 clusters with 286877 cMITEs (20.1%) implied genomes of the family Enterobacteriaceae, prone to multiple HGT events accordingly with the bibliography (69–71). Finally, a total of 5053 cMITEs (0.35%, 124 clusters) could also be removed from the final count as they were detected in metagenome-assembled genomes (MAGs) with <90% of completeness and/or >5% contamination (accordingly with the standards suggested by (72). A clustering analysis using stricter parameters, 100% id, although reduced the number to 592 mix-clusters (19995 cMITEs, data not shown), showed the persistence of mix-clusters with cMITEs shared among a wide taxonomic rank.

**Figure 5.**
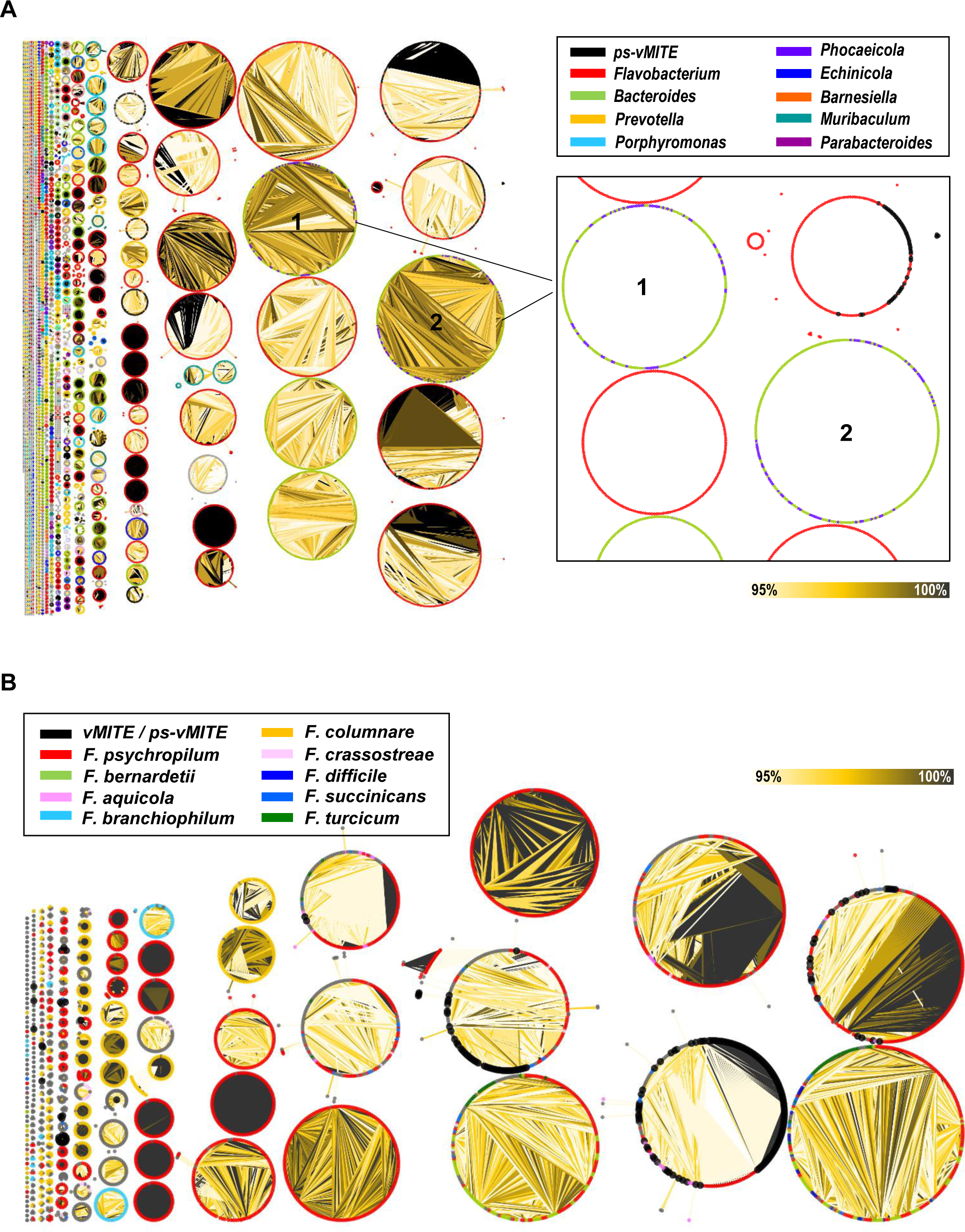
(A) Relation among the cMITEs found in Bacteroidetes genomes and the associated vMITEs (including *ps*-v-MITEs). Inlet is amplified the mix-clusters found within *Bacteroides* and *Phocaeicola* genera. (B) Net of cMITEs and vMITEs of Flavobacteriales. Nets were constructed with SSnet (cut-off: 95% and 99% coverage) and represented with Cytoscape (v. 3.10.1).

**Figure 6.**
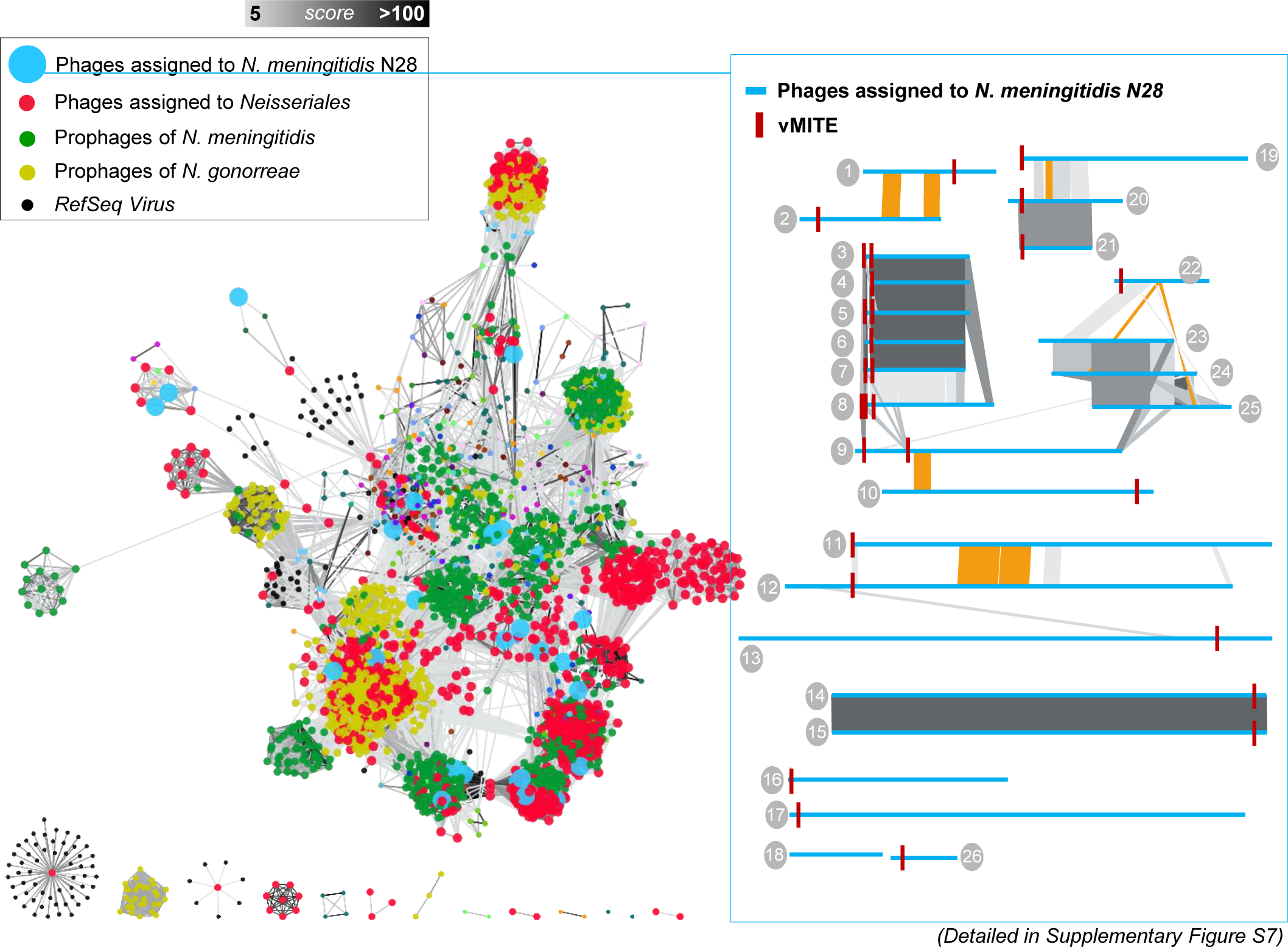
Relationships among the RefSeq virus and the 943 viral sequences associated to Neisseriales through MITEs sequences. Prophages already described for *N. meningitides* and *N. gonorreae* in (75) were included in the analysis. Net was constructed with Vconcat2 and represented in Cytoscape (v. 3.10.1). Only those relations with a score >5 were considered. Relationship among the viral sequences assigned to *N. meningititis* N28 is shown in the outlet of the figure at the right (see methods for accession numbers and a complete detail can be found in Supplementary Figure S7).

### MITEs in Virus, vMITEs and *ps*-vMITEs

Next, a large-scale systematic survey for viral MITEs (vMITEs) was performed. For this purpose, the NCBI viral database and the high-confidence-contigs of IMG/VR were used (see methods). Results showed that only 0.01% of the viral genome (complete or partial) were positive for vMITEs, with 671 viral sequences containing 1726 vMITEs (645 unique) (Supplementary File S3). Consistently with the low numbers of vMITEs detected, Zang et al (2018) also observed low number of vMITEs (0.2% of viral sequences from a total of 5170 viruses analyzed). Among them, more than one third (551 vMITEs, 31.9%) belonged to eukaryote viruses and 18 of them were previously described by Zhang et al., (2018); i.e, *Pandoravirus*, *Pithosvirus sibericum P1084-T* or the *Emilianaia huxleyi virus 156*. A total of 284 vMITEs were identified in already described bacterial phages such as phages of *Acidobacteriae UBA7540* (n=99)*, Prevotella* (n=41)*, Desulfobaccales* (n=25)*, Enterococcus faecium* (n=9)*, Kingella kingae*, *E. coli*, *Klebsiella pneumoniae, Pantoea stewartii* and *Pseudomonas cannabina* (Supplementary Table S5). However, for a total of 212 viral sequences, the host was unknown.

Given the low percentage of viruses containing vMITEs, it was questioned whether the high mutational rates of virus might lead ancestral vMITEs to have accrued mutations in the TIR or the TSD regions that masked their identification by bioinformatic programs. Those degenerated vMITEs, henceforth referred to as *pseudo*-vMITEs (*ps*-vMITES), were searched using each previously identified cMITE as a BLASTN query against the viral datasets, but allowing a range of mutations (c*ut-off*: 95% identity, 100% length coverage), a value among the mismatches observed previously (22,53). This approximation will further allow to link a virus sequence to its bacterial/archaeal host. This analysis revealed 5235 new *ps*-vMITES within 1840 viral sequences, 379 of them with an unidentified host (a higher number could be obtained with a less stricter criteria of 95% id in 70% of the length, 13608 *ps*-vMITEs in 11863 viral contigs, data not shown). Whether the degenerated *ps*-vMITEs are or not functional is unknown and further investigations must be addressed in this regard. Accounting both the vMITEs and the *ps*-vMITEs, the total number of these MGEs was of 6961, identified in 2798 viral sequences (Table 2). The average number of vMITEs (including *ps*-vMITEs) per sequence was of 2.74 (0.18 vMITE/kb), however, it is important to note that many of the viral sequences used may be incomplete. Majority of viral sequences contained two MITEs (59.7%), but up to 30 different vMITEs were found within a single viral sequence associated with *N. meningitidis* (IMGVR_UViG_2537561912_000003).

### Global clustering, connecting virus and host through their MITEs

The next step was to link the viral sequences (n=2510), with their putative host. All the previously identified MITEs (cMITEs, vMITEs and the *ps*-vMITEs) were clustered together and those groups containing vMITEs (vMITEs or *ps*-vMITEs) together cMITEs were closely examined (*cut-off*: 95% id, 100% length coverage). This allowed the association of 1880 viral sequences to a putative bacterial or archaeal host (Table 2 and Supplementary Table S6A-B). These associations were compared also with those resulting from iPHoP program (73) and resulted that 71.7% of them were paired with identical results than with MITEs (rest was unassigned by iPHoP). A total of 495 viral sequences had similar vMITEs than those cMITES found in a single prokaryotic genome, 586 to cMITEs present in several genomes from the same species, and another 711 viral sequences to cMITEs from several species from the same bacterial genus. Working as positive controls, 1501 of these associations were unambiguously validated as they were known virus-host pairs because they were isolated virus or through CRISPR assignments. A total of 379 new assignations were defined, viral sequences which putatively belong to *Gemmiger* phages (33), *Pseudomonas* (22), *Prevotella* (17), *Escherichia* (17), *Blautia* (13), *Neisseria* (12), *Flavobacterium* (12) and *Bacteroides* (11). Yet, 127 clusters trespass genus boundaries (97 clusters contained cMITEs from microbial genomes of the same family, 16 from the same order and 14 from class or phylum). Some of these exceptions implied microbial species with already frequent HGT events described; for example, one cluster shared vMITEs with those cMITEs from *Paenibacillus* (Bacillota) and *Chryseobacterium* (Bacteroidota) genomes, both genera being part of lignocellulose-degrading microbial consortia and sharing multiple MGEs (74). Similarly, 21 groups contained vMITEs with cMITEs from different genus of the Enterobacteriaceae family (*Escherichia, Klebsiella*, and *Enterobacter*, among others) with multiple HGT and sharing common MGEs as previously cited.

The significance of certain microbial groups with genomes enriched in cMITEs, prompted us to conduct a more thorough investigation using all available genomes (not only the RefSeq) in Bacteroidota and Neisseriales groups as examples to evaluate the discovery of new phages as Crassvirales (they infect Bacteroidetes bacteria, a prevalent group in the human gut microbiome) or increase the number of virus known in pathogenic *Neisseria*. Briefly, a total of 5837 genomes of Neisseriales were screened and new cMITEs were detected, increasing from 1639 to 6769 unique cMITEs within the Neisseriales group (Supplementary Table S8) which allowed to recruit 943 viral sequences from the viral databases (223 were previously identified). A total of 533 belonged to phage that putatively infect *N. meningitidis,* 296 to *N. gonorrhoeae* and 25 to *Kingella kingae*. Their relation with the Ref-viral sequences and also the *N. gonorrhoeae* prophages recently described (75) is showed in Figure 7. Although some clusters may be discerned, most of them are interconnected due to the numerous interchangeable pro-phages and HGT events among this genus. A total of 701 sequences presented an ANI over 95% previous published prophages, but novel clusters also appears. Importantly, the same vMITE sequence can be found in different viral sequences (different clusters) as shown, for example, in the putative phages of *N. meningitides* N28 genome (caption in Figure 7 and Supplementary Figure S7). In this case, the most likely scenario involves the transfer direction starting from *Neisseria* to the virus, followed by the spread of the MITEs within the phage genome. Other similar cases of different viral sequences with equal vMITEs were found for *Enterococcus* and *Bacteroides* phages (Supplementary Table 6). Regarding the Bacteroidota group, the new screening of 46051 Bacteroidota genomes resulted in an increased from 6122 to 81380 unique cMITEs and 929 viral sequences were finally localized (125 were previously identified using only the RefSeq). From them, 530 were unknown and assigned to phages of *Bacteroides* (204), *Prevotella* (83), *Flavobacterium* (66) and *Parabacteroides* (11). A net reconstruction showed our sequences not to be related with any of the crAssphages previously described; however, novel clusters were discovered (Figure 8). The recruitment in four fecal metaviromes from healthy human adults previously published, showed that some of them where highly represented (Supplementary Table S10).

**Figure 7.**
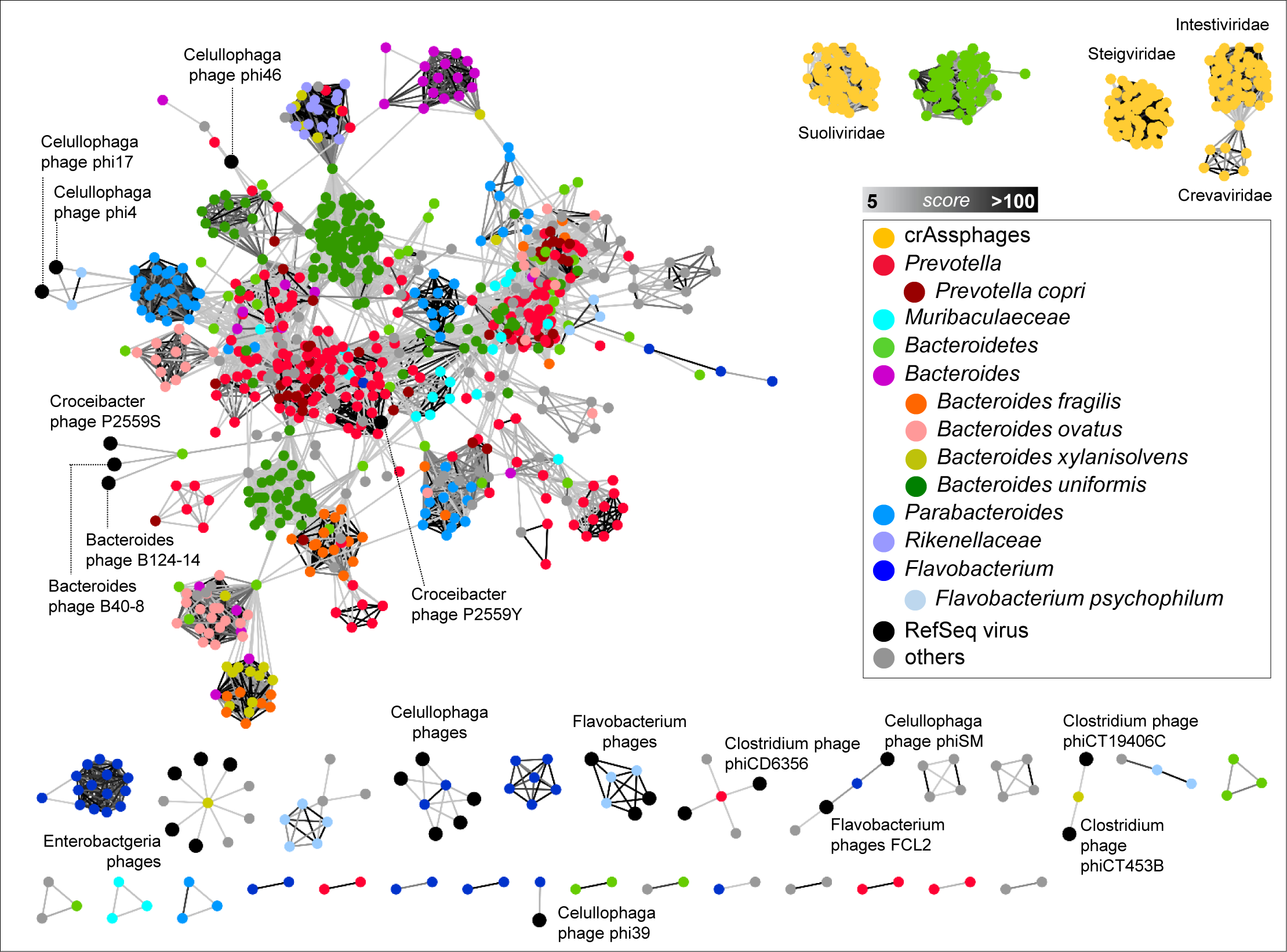
Relationships among the RefSeq virus and viral sequences associated to Bacteroidetes through MITEs sequences. Crassvirales already published in the NCBI database was also used in the analysis. Net was constructed with Vconcat2 and represented in Cytoscape (v. 3.10.1). Only those relations with a score >5 were considered.

In general, little is known about the exact mechanisms of cMITEs transfer, but even less, between virus and their host. However, some insights might be gleaned. One logical condition that would favor the mobility of MITEs between hosts and viruses is lysogeny. As prophages become integrated into the host genomes, evolution along time would increase the likelihood of a genetic interchange. Therefore, the 2798 viral sequences with a vMITE were screened to determine their lysogenic or lytic nature. Despite that many of the environmental sequences may be incomplete, 8.8% of them were identified as putative prophages (Supplementary Table S7), a slightly higher than the 5.5% of the viral sequences from the complete database used.

Another important need to MITE mobility is an external transposase. In this regard, several IS-transposases sharing equal (or similar) TIR domain than the cMITEs have been described (22,53,76,77). Hence, viruses with their own transposases might be used also independently from the host genome. Transposases were screened within the viral sequences with vMITEs and 864 (34.4%) were found positive. In this regard, further experiments would be required to clarify if TIR sequences are the only condition to allow a MITE to be mobilize or other requirements are needed.

## DISCUSSION

### Census of cMITEs

A census of cMITEs across the RefSeq prokaryotic genomes was conducted, unveiling their presence in more than half of the screened sequences (58.4%) and encompassing thousands of unique cMITEs. In contrast to eukaryotes, where cMITEs have been extensively described and considered essential in the evolution of their genomes (33), the majority of cMITEs in prokaryotic genomes have predominantly gone unnoticed due to their intergenic position and unknown functional roles. In general, cMITEs are considered selfish elements that may impose a considerable cost on cells leading to diverse fitness outcomes (27,78). However, the ‘permitted’ expansion of some cMITEs in specific prokaryotic genera and their persistence throughout evolution in specific *loci* (45,53), suggests a broader significance beyond a mere self-serving behavior and, instead, benefits from these “DNA-parasites” may be gained by the host over the long-term course of their co-evolution (79).

cMITEs were widely distributed across Bacteria and Archaea but, as showed by the standard deviation observed for some genus, very heterogeneously (Supplementary Table 1). Meanwhile they appeared amplified extraordinarily in some groups, they were mostly absent in genera where thousands of genomes are available (Figure 2). For example, cMITEs were detected in hundreds of copies in *Nostocales*, *Oscillatoriales* or *Leptospirales* genera, or present in hundreds of genomes of clinical pathogens such as *Neisseria, Streptococcus, Mycobacterium, Legionella, Salmonella* or *Shigella*. On the contrary, they were almost absence in *Staphylococcus*, *Klebsiella*, *Acinetobacter*, *Enteroccoccus* or *Mycobacteroides*, despite the description of several MGEs among their genomes, particularly those related with antibiotic resistance gene transmission, (*80,81*). While intricate microbial defense systems that regulate or eliminate MGEs have been extensively documented (19), limited knowledge exists regarding the persistence of MITEs and the existence of well-conserved purging strategies specifically controlling cMITEs’ proliferation throughout the genome

Although different strategies might be followed by cMITEs to expand within a cell genome, one plausible explanation would be polyploidy. The existence of multiple DNA replicons in the cell cytoplasm at once, would enhance the possibility to be copied and integrated in several *loci* along the genome. This would explain, for example, the burst of cMITEs found in some Cyanobacteria genomes (which also have a high number of DNA repetitions observed, (82,83)), or also, some cases of Spirochaetes and Neisseriales (84,85). Other cases, like the endosymbiont *Wolbachia,* with the highest density rate of cMITEs detected, it might possible that the large number of IS elements present in this genera would promote their mobilization (86). In summary, the prevalence of multiple cMITEs in most of the genomes of a particular genus will respond to a mechanism where, once that a cMITE is established in the genome, its expansion to similar strains and in substantial numbers (media was of 9 cMITEs/genome) becomes highly probable, especially in the absence of purging strategies against these MGEs. This trend is supported by the low percentage of the genomes that had one single cMITE, 2.5%, and because for about 30% of genera, the cMITEs where present in all the species genomes sequenced (Supplementary Figure S5). The low number of observed MITEs in some genomes may be also attributed to a limited genome-flexibility; e.g., the absent of cMITEs in free-living microbes with streamlined genomes, such as *Pelagibacteres*. In those cases, the pressure to maintain optimized the number of genes (functionality) in such short genomes avoids the inclusion of foreign DNA. Indeed, although *Pelagibacter* genomes encode DNA uptake genes, no horizontally transferred DNA has been detected, nor do they even contain any known transposable elements (19,87,88).

Along this lines, other interesting observation was that, in genera such as *Pseudomonas* or *Vibrio,* with hundreds of species with a high variety of niche occupations, cMITEs were found in only 60% of their genomes, indicating a substantial influence of environment in the cMITEs maintenance. However, the number of unique cMITEs surpass three or four orders of magnitude the observed media, 77.46 *versus* the 19309 different MITEs found in *Pseudomonas* and 2361 in *Vibrio* (Figure 2). Furthermore, the circle nets of *Pseudomonas* (Figure 3A), and in a lesser extent those of *Vibrio*, showed a higher number of outsider “spikes”. One potential scenario is a heightened mutational bias acting against these elements, leading to an increased number of divergent cMITEs. Alternatively, the cMITE repertoire may expand to boost the fitness of their hosts in diverse environments, facilitating mobilization or gene activation. This, in turn, could induce beneficial rearrangements or contribute to the enrichment of the host’s gene pool. Regarding the little number of vMITEs found in virus (including *ps*-vMITEs), accounting less than 0.005% of the examined viral sequences, it is very probable that the small viral genome size, their high gene density, and the physical constraints of genome size-virus per particle, would limit the incorporation of new insertions into the viral genome and, unless this confers a clear beneficial advantage (or neutral), active elimination through counterselection will happen.

### Specificity of cMITEs as a tool for pairing microbial host with viruses

In sharp contrast with the conventional view of MGEs, cMITE exhibit a relatively limited mobility. Clustering analysis of the detected cMITEs revealed that, while 2.6% of these elements are dispersed among genomes of different families, orders or classes; the vast majority of cMITEs, 97.4%, ‘move’ confined within the boundaries of genera. Importantly, 23% of them were specific within a single microbial species. The exceptions may be explained in the context of HGT events among distant taxa, as it has been described multiple times in literature (71,89). The mobility of cMITEs is well-documented within the same genome or among closely related species genomes (43,45,53,54); but little is known about their flow across different taxonomic ranks and more experiments are needed in this regard. cMITEs are characterized by the need the activity of an outsider transposases to be “copy-pasted” among sequences; then, cMITEs sharing only the same TIR (with different internal sequence), could be mobilized by several transposases across very different species as happens with other MGEs (90,91). However, it is puzzling that cMITEs were mainly conserved only beyond the genus limits (non inter connected circles in the nets of Figure 3). This barrier suggests the existence of restrictive cellular mechanisms against their dispersion, and it is plausible that, to be embraced in the new genome host, the internal sequence of the cMITE must meet additional genomic features, such as an appropriate GC-content (supported by the correlation found among the GC-content of the cMITE and the host genome) or even a concrete methylation pattern. Interestingly, the existence of equal cMITE sequences in different species might work as hallmarks of HGT events among them and tracking the cMITEs sequences along, might be a useful tool to understand the gene flow among closely cells or even symbiosis; i.e., the common cMITEs among the spider *Oedothroax gibbosus* and its *Cardinium* endosymbiont (92)).

In this work, the global clustering of all the microbial cMITEs detected and their viral counterparts, vMITEs (*ps*-vMITEs included), allowed to pair up to 51655 host sequences to 1880 viral sequences (Supplementary Table S6). At least 1501 (79.8%) cases were unambiguously assigned as they belong to previously described cases, confirming then that the existence of a MITE in a virus sequence would allow to determine, at least, the genus of the host, if not the specie. This way, 379 new viral sequences were associated to *Pseudomonas*, *Ruminococcus*, *Faecalibacterium* or *Prevotella*, among others. Particular interesting was the expansion of the Neisseriales and Bacteroidota virospheres and further analysis of some of the new clusters is worthwhile.

Recognition by most viruses to their hosts is highly specific and instances of phages infecting members of different genera are rare (93,94). However, due to the small number of sequences in viral genomes, it seems most likely that frequent HGT events among closely related species contribute more extensively to the cMITE distribution in prokaryotic cells. Certainly, transduction facilitated by virus might also significantly contribute to the dissemination of these cMITEs. From the cell genome, and facilitated by a prophage form, the cMITE may be copied and integrated into the viral genome. While the transfer may be initially purely stochastic, subsequent purging and fitness evolution will contribute to removal or fixation of these MITEs in the genomes of the viruses. This scenario would explain, for example, the nearly identical matches of one single vMITE from one viral sequence to 21 cMITEs found in different *Pseudomonas* spp. Therefore, viruses would simply serve as vectors, albeit “bad vectors” (by the low number detected), for the horizontally transferred cMITEs.

Undiscovered viruses beckon researchers, and a vast reservoir of viral sequences lingers within databases, largely untapped, primarily due to several challenges, including those associated with the identification of their infection targets. We have shown that cMITEś genus specificity could be a useful tool to bridge these gaps, especially when dealing with novel viral sequences for which no other tools are available beyond their assembled sequence. Due to time and computational constraints, this study has been limited to publicly available bacterial RefSeqs and the archaeal genomes. However, the discovery of new cMITEs is possible when leveraging all genomes within specific genera and revealing novel virus associations as showed with the Bacteroidota and Neisseriales groups. Certainly, exploring MITEs in metagenomes and metaviromes will open up an extended landscape to unveil novel host-virus relationships.

## Acknowledgements

We thank Prof. J. Antón for continuous support of this research project.

## Author contributions

FNM, ABMC, RR, SGJ and ACL performed bioinformatic analysis. ABMC planned experiments and wrote the manuscript.

## SUPPLEMENTARY DATA

### Supplementary Tables

**Supplementary Table S1.** Distribution of cMITEs found in genus with more than 20 sequenced genomes. Standard deviations, means and coefficient of variation are indicated.

**Supplementary Table S2.** List of *Staphyloccocus* genomes accessions with cMITEs detected.

**Supplementary Table S3.** List of prokaryotic genomes with higher number of cMITEs. Only the 100 first genomes are listed.

**Supplementary Table S4.** Archaeal and bacterial genomes ordered by number MITEs/Mb.

**Supplementary Table S5.** List of viral sequences containing vMITEs, their NCBI-taxonomy and their assigned host.

**Supplementary Table S6. (A)** Virus-Host clusters associations in archaeal genomes. **(B)** Phage-Host clusters associations in bacterial genomes.

**Supplementary Table S7.** CheckV results for viral sequences with vMITEs or *ps*-vMITEs.

**Supplementary Table S8.** Viral sequences with similar cMITEs than found in Neisseriales genomes.

**Supplementary Table S9.** Viral sequences with similar cMITEs than found in Bacteroidetes genomes.

**Supplementary Table S10.** Recruitment (rpkg) of Bacteroidota phages assigned through MITEs in healthy adult fecal human metaviromes. Sequences collections were downloaded from the following projects; PRJNA491626 (SRR7892428 and SRR7892426) (Yinda, Vanhulle et al. 2019); PRJEB9524 (ERR906942 and ERR906934) (Monaco, Gootenberg et al. 2016) and PRJNA318788 (SRR3403840) (Chehoud, Dryga et al. 2016). Crassphages already published were also included in the analysis (in green).

### SUPPLEMENTARY FIGURES

**Supplementary Figure S1.** (A) cMITEs distribution in relation with the prokaryotic genome size in Bacteria and (B) for Archaea. Bars (right *y axe*) indicate the number of genomes analyzed. (C) Distribution of the unique cMITEs in Bacteria and (D) in Archaea.

**Supplementary Figure S2.** (A) Box plots indicates the number of cMITEs in relation with the GC content of the host genome. Grey bars (right *y axe*) indicate the distribution of the genomes accordingly with their GC-percentage. (B) Distribution of the GC-content of the cMITE and the GC-content of the host genome. (*P. corr, Pearson Correlation).

**Supplementary Figure S3.** Distribution of the number of cMITEs found in Bacteria (A) and Archaea (B) genomes accordingly with the coding density of the genomes that host them. Grey Bars (right *y axe*) indicate the number accordingly with their coding density. Colors of the dot respond to their phylum taxonomy.

**Supplementary Figure S4.** Intergenic and intragenic positions of MITEs in Archaea, Bacteria and the viral sequences from GenBank and JGI IMG/VR databases.

**Supplementary Figure S5.** Relation of number of genomes within a specie and genus that contains cMITEs. For example: 28.7% of the strains of a single species presented cMITEs in all their genomes (100%), meanwhile, for 30.49% of the strains for other species, presented cMITEs in only 10% of the genomes.

**Supplementary Figure S6.** Distribution of genomes with MITEs in Cyanobacteria phylum (A) and Stenosarchaea group (B) genomes, accordingly with size of genome (right *y axis*) and number of MITEs/genome (*x axe*). Color respond to last taxon classified. Number of genomes in each range of number of MITEs is marked with left *y axis.*

**Supplementary Figure S7.** Comparison of *Neisseria meningitides* new phage sequences (see methods for accession sequences). Despite the vMITEs shared, phage sequences were not related. Note that different putative phages shared the same *N. meningitidis* cMITE, which contains a fragment of about 100 pb conserved in a transposase nearby (Fig. S9B).

### SUPPLEMENTARY FIGURES

#### Files of sequences

**Supplementary File 1.** Sequences of cMITEs detected in Bacteria genomes (*fasta* format). The hosting microbial species and inferred NCBI-taxonomy are indicated in the name of each sequence.

**Supplementary File 2.** Sequences of cMITEs detected in the Archaea genomes (*fasta* format). The hosting microbial species and inferred NCBI-taxonomy are indicated in the name of each sequence.

**Supplementary File 3.** Sequences of vMITEs detected in the virus sequences from the NCBI and IMG/VR v.4.1 database (*fasta* format). Virus, microbial host (if known) and inferred NCBI-taxonomy is stated in the name of each sequence.

**Supplementary File 4.** Sequences of *ps*-vMITEs detected in the virus sequences from the NCBI and IMG/VR v.4.1 database (*fasta* format). Virus, microbial host (if known) and inferred NCBI-taxonomy is stated in the name of each sequence.

**Supplementary File 5.** Sequences of cMITEs obtained from 5837 genomes of Neisseriales.

**Supplementary File 6.** Sequences of cMITEs obtained from 46051 genomes of Bacteroidota.

