## Supplementary Figures for "Bridging Viruses and Prokaryotic Host through Miniature Inverted-repeat Transposable Elements (MITEs)"

**Supplementary Fig. S1.** Nadal-Molero *et al.* 2024

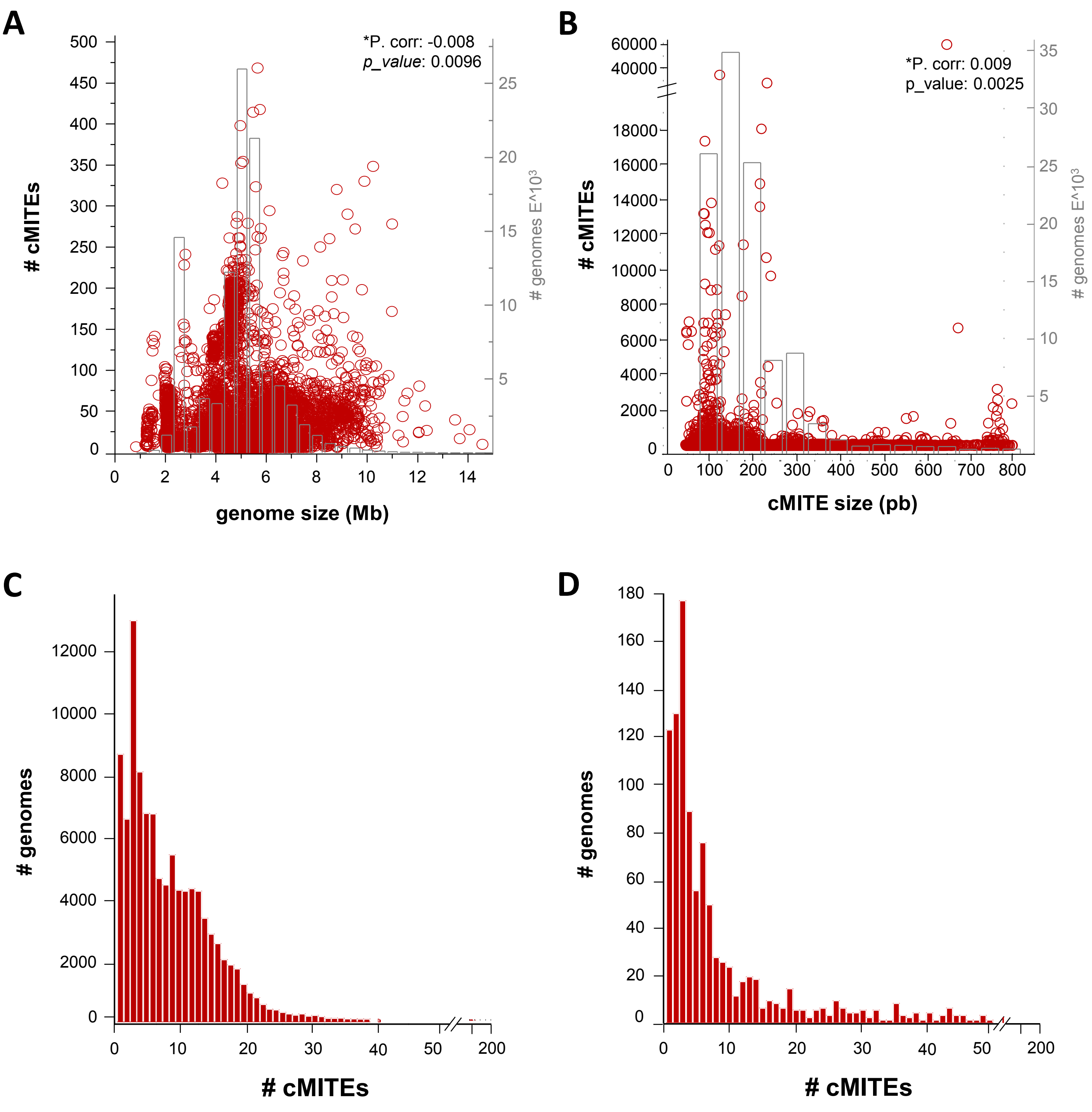


**Supplementary Figure S1**. (A) Distribution of cMITEs in relation to the prokaryotic genome size in Bacteria and (B) for Archaea. Bars (right *y*-axis) indicate the number of genomes analyzed. (C) Distribution of unique cMITEs per genome in Bacteria and (D) in Archaea.

**Supplementary Figure S2.** Nadal-Molero *et al.* 2024

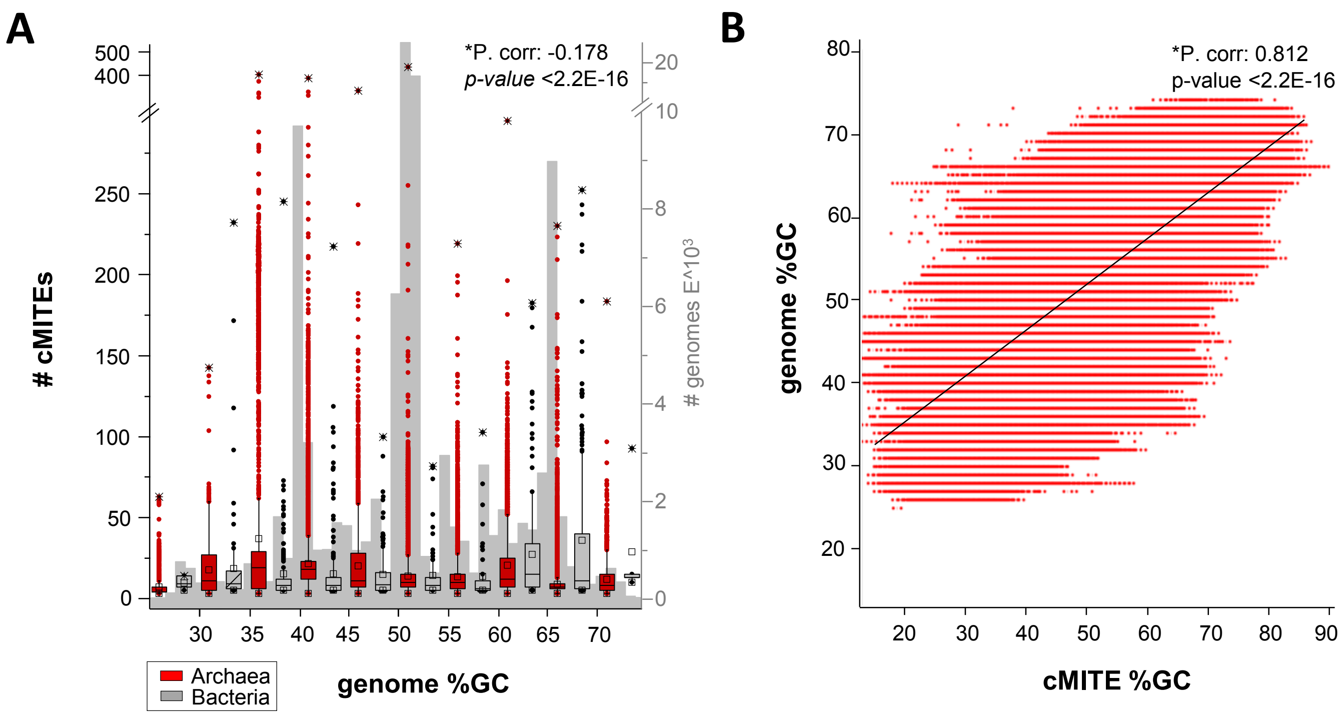


**Supplementary Figure S2. .** (A) Box plots indicate the number of cMITEs in relation to the GC content of the host genome. Grey bars (right *y*-axis) indicate the distribution of the genomes according to their GC percentage. (B) Distribution of the GC-content of the cMITE and the GC-content of the host genome. (*P. corr, Pearson Correlation).

**Supplementary Figure S3.** Nadal-Molero *et al.* 2024

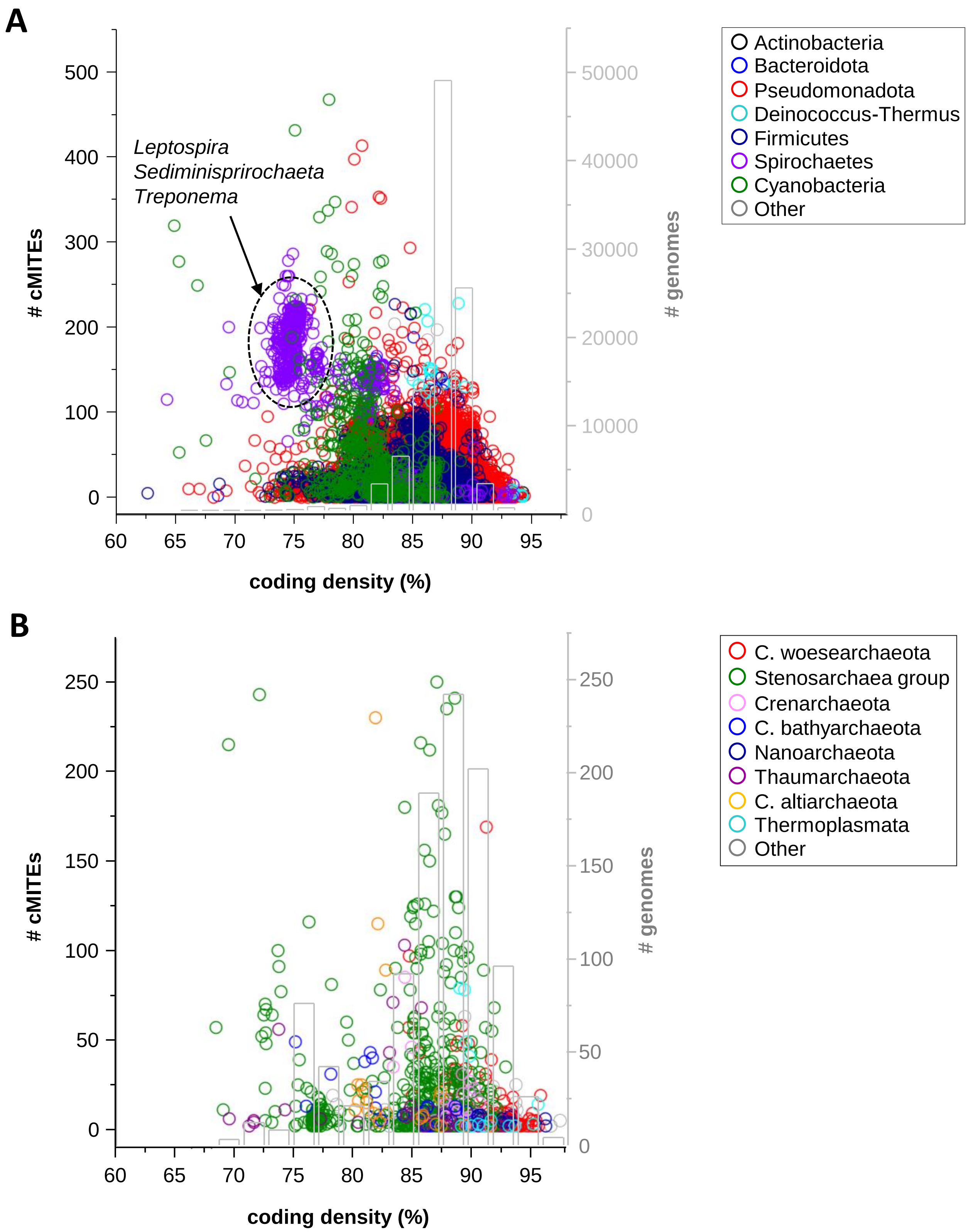


**Supplementary Figure S3**. . Distribution of the number of cMITEs found in Bacteria (A) and Archaea genomes (B) based on the coding density of the hosting genomes. Grey bars (right *y*-axis) indicate the number corresponding to their coding density. Dot colors represent their phylum taxonomy.

**Supplementary Figure S4.** Nadal-Molero *et al.* 2024

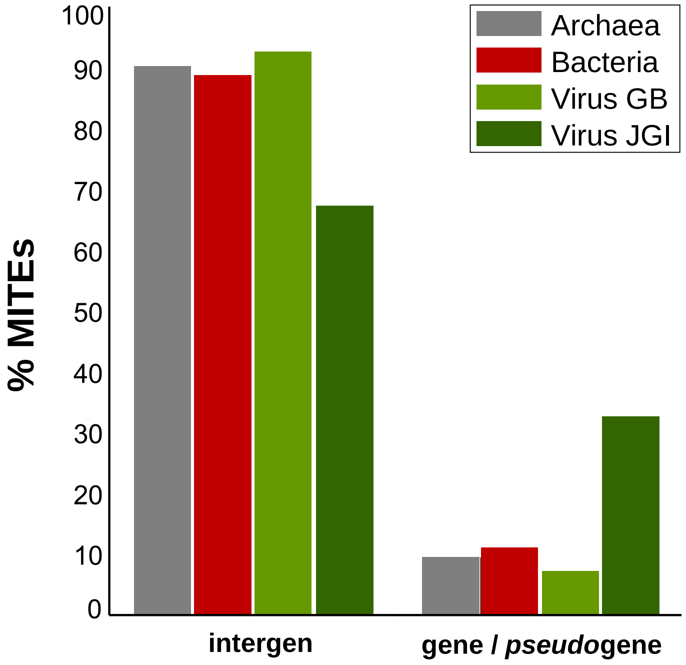


**Supplementary Figure S4.** Distribution of intergenic and intragenic positions of MITEs in Archaea, Bacteria and viral sequences from GenBank and the JGI IMG/VR databases.

**Supplementary Figure S5.** Nadal-Molero *et al.* 2024

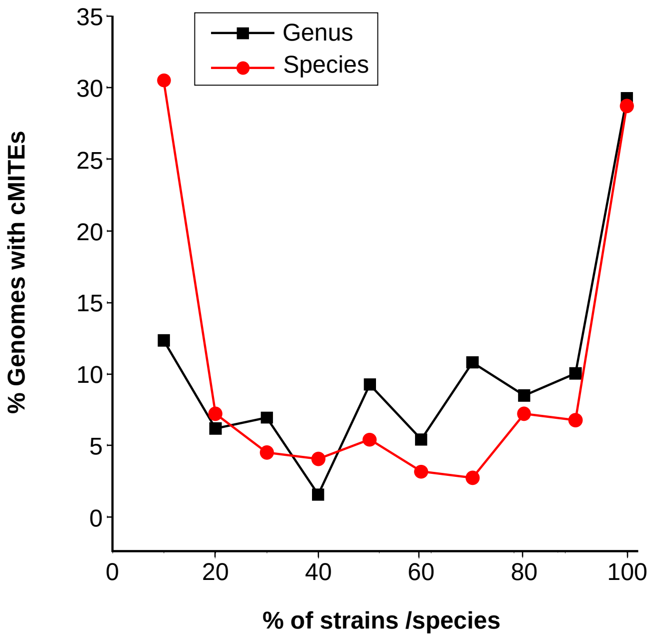


**Supplementary Figure S5.** Relationship between the number of genomes within a species and genus that contains cMITEs.

**Supplementary Figure S6**. Nadal-Molero *et al.* 2024

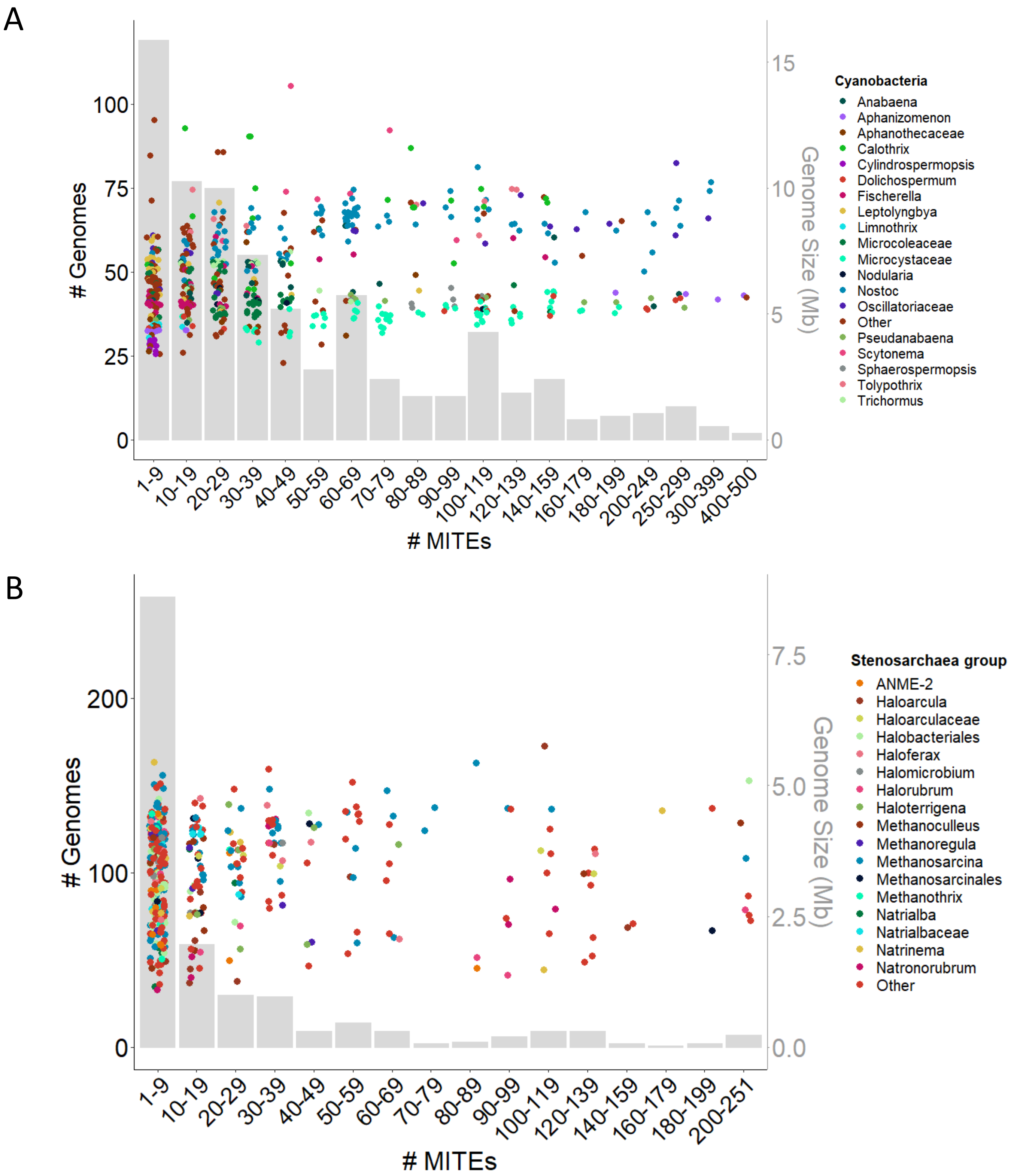


**Supplementary Figure S6.** Distribution of genomes with MITEs in the Cyanobacteria phylum (A) and Stenosarchaea group (B) genomes, according to the size of the genome (right *y*-axis) and the number of MITEs/genome (*x*-axis). Colors correspond to the last taxon classified. The number of genomes in each range of the number of MITEs is marked with left *y axis.*

### **Supplementary Figure S7**. Nadal *et al.* 2024

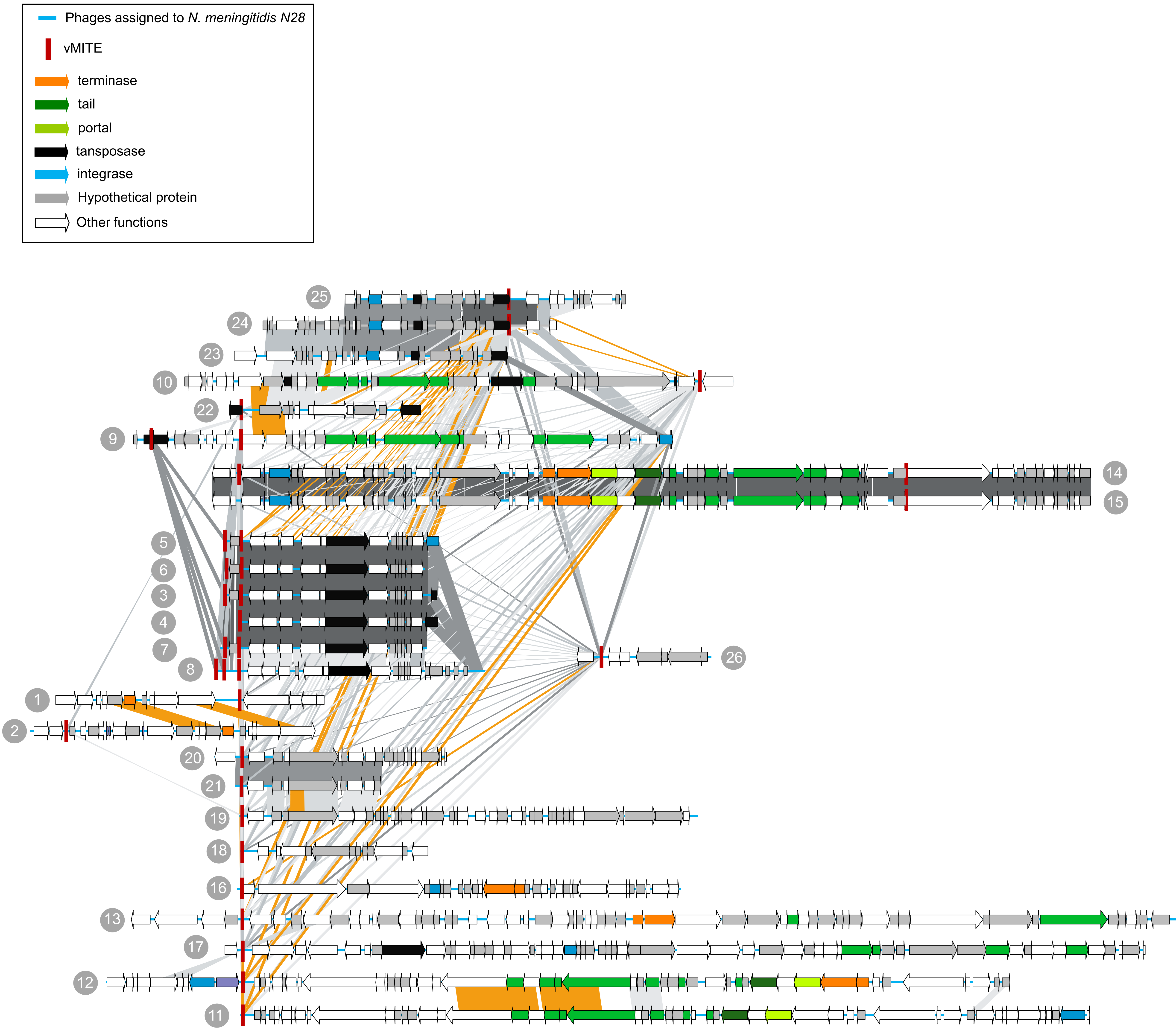


**Supplementary Figure S7.** Comparison new phage sequences assigned to *Neisseria meningitides* N28 (see Materials and Methods for accession sequences). Note that despite the shared vMITEs, phage sequences were not always related. Different putative phages shared the same *N. meningitidis* cMITE, which contains a fragment of about 100 pb conserved in a transposase nearby.
